# Galantamine Attenuates Autoinflammation in a Mouse Model of Familial Mediterranean Fever

**DOI:** 10.1101/2022.06.10.495119

**Authors:** Ibrahim T. Mughrabi, Mahendar Ochani, Mirza Tanovic, Ping Wang, Betty Diamond, Barbara Sherry, Valentin A. Pavlov, Seza Ozen, Daniel L. Kastner, Jae Jin Chae, Yousef Al-Abed

## Abstract

Autoinflammatory diseases, a diverse group of inherited conditions characterized by excessive innate immune activation, have limited therapeutic options. Neuroimmune circuits of the inflammatory reflex control innate immune overactivation and can be stimulated to treat disease using the acetylcholinesterase inhibitor galantamine. Here, we tested the efficacy of galantamine in a rodent model of the prototypical autoinflammatory disease familial Mediterranean fever (FMF). Long-term treatment with galantamine attenuated the associated splenomegaly, amyloidosis, and anemia that are characteristic features of this disease. Further, treatment reduced inflammatory cell infiltration into affected organs and a subcutaneous air pouch. These findings suggest that galantamine attenuates chronic inflammation in this mouse model of FMF. Further research is warranted to explore the therapeutic potential of galantamine in FMF and other autoinflammatory diseases.

## Introduction

The autoinflammatory diseases are a group of inherited conditions characterized by fever and antigen-independent systemic inflammation. A subset of these conditions are caused by mutations in inflammasome proteins leading to dysregulated IL-1β and IL-18 production [1]. The prototypic and most common example of these disorders is familial Mediterranean fever (FMF), found almost exclusively in the eastern Mediterranean basin, with a prevalence as high as 4% in some populations [2]. FMF is a recessively inherited condition caused by gain-of-function mutations in *MEFV*, which encodes the inflammasome protein pyrin [3]. Due to consequent elevations in pro-inflammatory cytokines, patients experience periodic fevers and painful inflammatory attacks that affect the skin, joints and body cavities. The inflammatory episodes are accompanied by large infiltrates of polymorphonuclear leukocytes in affected tissues with a concomitant increase in acute phase reactants [4]. FMF is a debilitating and incurable disease that can lead to death due to amyloidosis and organ failure. The standard treatment for FMF is lifelong administration of colchicine, which controls the febrile attacks in two-thirds of patients and prevents lethal amyloidosis. However, in about one-third of patients, colchicine is inadequate and can be toxic at high doses [5]. In those patients, newer biologic agents, such as anakinra and canakinumab, have shown efficacy in treating the recurrent attacks [6,7]; however, their high cost, regional unavailability, and adverse effects limit their clinical use. Therefore, there is a need for novel and safe therapies that can control FMF inflammation and prevent its chronic complications. Such therapies might also be useful in other autoinflammatory and innate immune spectrum diseases, such as gout, diabetes and heart disease.

Recent advances have revealed a neurophysiological mechanism that controls inflammation via the vagus nerve, termed the inflammatory reflex [8]. Numerous studies have explored this reflex as a potential therapy in various animal models of inflammatory and auto-immune disease, using electrical and pharmacological means to stimulate the vagus nerve [9–13]. These studies have demonstrated that signaling along the efferent arm of the inflammatory reflex controls pro-inflammatory cytokine release in experimental models of inflammation. This effect is brought about by the interaction of acetylcholine (ACh) with the alpha 7 nicotinic acetylcholine receptor (α7nAChR) expressed on the surface of macrophages and other immune cells [14]. These preclinical findings were carried forward in recent pilot clinical trials using electrical stimulation or centrally-acting cholinergic agents, such as galantamine, to activate the inflammatory reflex and successfully treat chronic inflammatory diseases. For example, implanted electrical vagus nerve stimulators improved symptoms in patients with rheumatoid arthritis and Crohn’s disease [15,16], and long-term pharmacologic treatment with galantamine alleviated inflammation and metabolic dysfunction in metabolic syndrome patients [17].

Modulating the inflammatory reflex is a clinically-tested approach to treat chronic inflammation. However, the therapeutic potential of this physiological mechanism has not been investigated in autoinflammatory syndromes, which currently have limited therapeutic options. Here, we utilize a mouse model of the most common autoinflammatory disease, FMF, to study the therapeutic potential of galantamine, a clinically approved drug for Alzheimer’s disease and a pharmacologic activator of the inflammatory reflex [18]. Our findings demonstrate that long-term treatment with galantamine attenuates several markers of inflammation and chronic disease complications in an FMF-knock in (FMF-KI) mouse model that harbors a human FMF mutation. Additionally, the persistence of the effect while blocking components of the inflammatory reflex suggests the engagement of other neuroimmune circuits in this chronic inflammatory model which warrant further exploration.

## Materials and Methods

### Reagents and chemicals

Ultra-pure LPS was purchased from InvivoGen (San Diego, CA). Galantamine, huperzine A, and colchicine were from Tocris Bioscience (Bristol, UK). Acetylcholine and nicotine were from Sigma Aldrich (St. Louis, MO).

### Mice

Heterozygous FMF-KI mice (*Mefv*^V726A/+^) were provided by Dr. Daniel Kastner (NHGRI, NIH) and bred in house to generate FMF-KI homozygotes (*Mefv*^V726A/V726A^) [19]. All experiments were performed on homozygous FMF-KI mice. α7nAChR-KO mice were provided by Dr. Kevin Tracey (Feinstein Institutes for Medical Research, NY) and were crossbred with FMF-KI mice to generate combined FMF-KI/α7nAChR-KO mice. Animal experiments were conducted according to the US National Institutes of Health guidelines and were approved by the Institutional Animal Care and Use Committee (IACUC) of the Feinstein Institute for Medical Research.

### Animal experiments

For long-term treatment experiments, 1.5-week-old male and female mice were treated with galantamine dissolved in PBS twice daily for ∼8 weeks. Galantamine was injected intraperitoneally (i.p.) at an escalated dose (4-12 mg/kg; increased 4 mg every 2 weeks) unless indicated otherwise. In some experiments, treatment started at age ∼4 weeks in mice with established disease and continued for ∼4 weeks escalating the dose every 2 days during the first week. The size of the spleen was used to assess response to long-term treatment since it represents the cumulative effects of ongoing systemic inflammation and immune cell expansion. To perform a complete blood count, mice were sacrificed at the indicated times and blood obtained by cardiac puncture. A 100 μL aliquot of blood was transferred to a tube containing 10 μL EDTA, incubated on ice for 30 minutes and then analyzed using a hematology analyzer according to manufacturer’s instructions. To obtain serum for cytokine assay, blood was left to clot for 2 hours at room temperature, then centrifuged at 2000 x g for 20 minutes and serum collected. Inflammatory markers were measured using V-Plex Proinflammatory Panel 1 (Mesoscale Discovery, Rockville, MD), and Serum Amyloid A ELISA Kit (R&D, Minneapolis, MN) following the manufacturer’s instructions.

### Air pouch model

The air pouch model was performed as previously described [20] with minor modifications. Mice received either galantamine (6-12 mg/kg, i.p.) or PBS twice daily over the week of air pouch generation. Dorsal air pouches were established in male and female FMF-KI mice (ages ∼8-14 weeks) by injecting 6 mL of air on days 0 and 3. On day 6, mice were euthanized and pouch contents were collected by injecting 3 ml of 2 mM EDTA in PBS and total cells counted using a hemocytometer.

### Vagotomy

After isoflurane anesthesia, ∼7-week-old animals were placed in the supine position and subjected to a 2-cm midline abdominal incision. The esophagus was then exposed just below the diaphragm where it joins the stomach by pulling the stomach downward. The anterior and posterior vagi were then visualized and cut, removing part of the nerve on each side to prevent regeneration. The spleen dimensions were measured before closing the wound. Animals were allowed to recover for a week before starting treatment with galantamine for 4 weeks (escalated to 12 mg/kg, twice daily). During the period of treatment, animals were kept on a semi-liquid diet to prevent stomach bloating.

### Cell Culture and inflammasome activation

Bone marrow cells were harvested from the tibiae and femora of male and female FMF-KI mice (ages 8 to 16 weeks) and differentiated for 7 days in Iscove’s Modified Dulbecco’s Media (IMDM) supplemented with 20 ng/ml of M-CSF (Peprotech, Rocky Hill, NJ), 10% heat-inactivated fetal bovine serum (FBS; Invitrogen, Carlsbad, CA), 1 mM sodium pyruvate, and 1% Pen/Strep (Invitrogen). Differentiated macrophages were then detached by scraping, counted, and re-plated in 12-well (1 × 10^6^ cells/well), 24-well (5 × 10^5^ cells/well), or 96-well (1 × 10^5^ cells/well) plates in RPMI (Thermo Fisher, Waltham, MA) supplemented with 10% FBS and 1% Pen/Strep and incubated overnight. The next day, medium was changed to Opti-MEM containing acetylcholine (100 μM), colchicine (3 μM), or nicotine (10-100 μM) for 30 minutes. Cells were then primed with LPS (1 μg/ml) for 6 hours. Supernatants were analyzed by IL-1β Duoset ELISA kit (R&D) following the manufacturer’s instructions.

### Western blot

Kidneys were homogenized on ice in T-PER (Thermo Fisher) containing Halt phosphatase and protease inhibitor cocktail (Thermo Fisher). Protein samples (50 μg) were then resolved on Novex 16% Tris-Glycine gel (Invitrogen) and transferred onto 0.22 μm PVDF membrane (BioRad) using a wet transfer system for 1 hour at 100 mV. Membranes were then incubated with primary antibody (anti-SAA A1, AF2948, 1:1000, R&D) overnight at 4 °C, washed the next day, and incubated with secondary antibody for 1 hour. Membranes were then stripped with Restore buffer (Thermo Fisher) for 15 minutes, then washed, blocked, and stained for actin as described.

### Flow cytometry

Single-cell suspensions from the spleen were obtained by flushing the organ with cold medium and expressing the cells by gentle compression. Peritoneal cells were collected by infusing the peritoneum with 5 mL cold PBS using a catheter needle then retrieving the lavage. Cells were stained with fluorophore-conjugated antibodies in staining buffer on ice in the dark for 30 minutes. Cells were washed with 2 ml FACS buffer, followed by 1ml PBS, then fixed by adding 240 μl of PBS and 60 μl of buffered formalin, after which data were acquired immediately. Anti-CD11b PE, anti-Ly6G FITC, anti-F4/80 Alexa647 antibodies were purchased from BD Bioscience (San Jose, CA) and used at a 1:60 dilution. Cells were analyzed using BD LSR II (BD Bioscience) and FlowJo software (Three Star, Ashland, CO).

### Statistical analysis

Statistical analyses were performed with Graph Pad Prism (Graph Pad Software Inc., La Jolla, CA). The significance level was set to α = 5% for all comparisons. Mann-Whitney *U* test was used to compare two groups. For *in vitro* experiments, one-factor ANOVA with post hoc Tukey’s multiple comparison test was used to compare multiple groups. Data are expressed as mean ± SEM unless indicated otherwise.

## Results

### Long-term treatment with galantamine prevents splenomegaly in FMF-KI mice

FMF-KI mice are characterized by severe local and systemic inflammation, massive splenomegaly, joint swelling, and weight and hair loss. To examine the role of the inflammatory reflex in autoinflammation, we studied the long-term effect of galantamine treatment on gross disease features in these animals. Treatment of 1.5-week-old FMF-KI mice with galantamine twice daily for 8 weeks significantly reduced the size of the spleen compared with PBS-treated controls (Fig 1 *A* & *B*). The long-term treatment had no apparent toxic effects, except for a small, but significant, loss in whole body weight (Fig 1 *C*), which is a previously reported side effect of the drug [21,22]. To determine whether galantamine’s effect on splenomegaly is mediated by its acetylcholinesterase (AChE) inhibitory activity, we treated FMF-KI mice with huperzine A, a potent centrally- and peripherally-acting AchE inhibitor that is structurally different from galantamine. After 4 weeks of treatment with huperzine A, we observed a comparable decrease in spleen size (Fig 1 *B*), which indicates that galantamine’s actions on splenomegaly are likely mediated by increased cholinergic activity achieved by AchE inhibition. Together, these findings demonstrate that long-term treatment with galantamine attenuates some of the physical signs of disease in FMF mice, possibly by potentiating cholinergic signaling.

**Figure 1.**
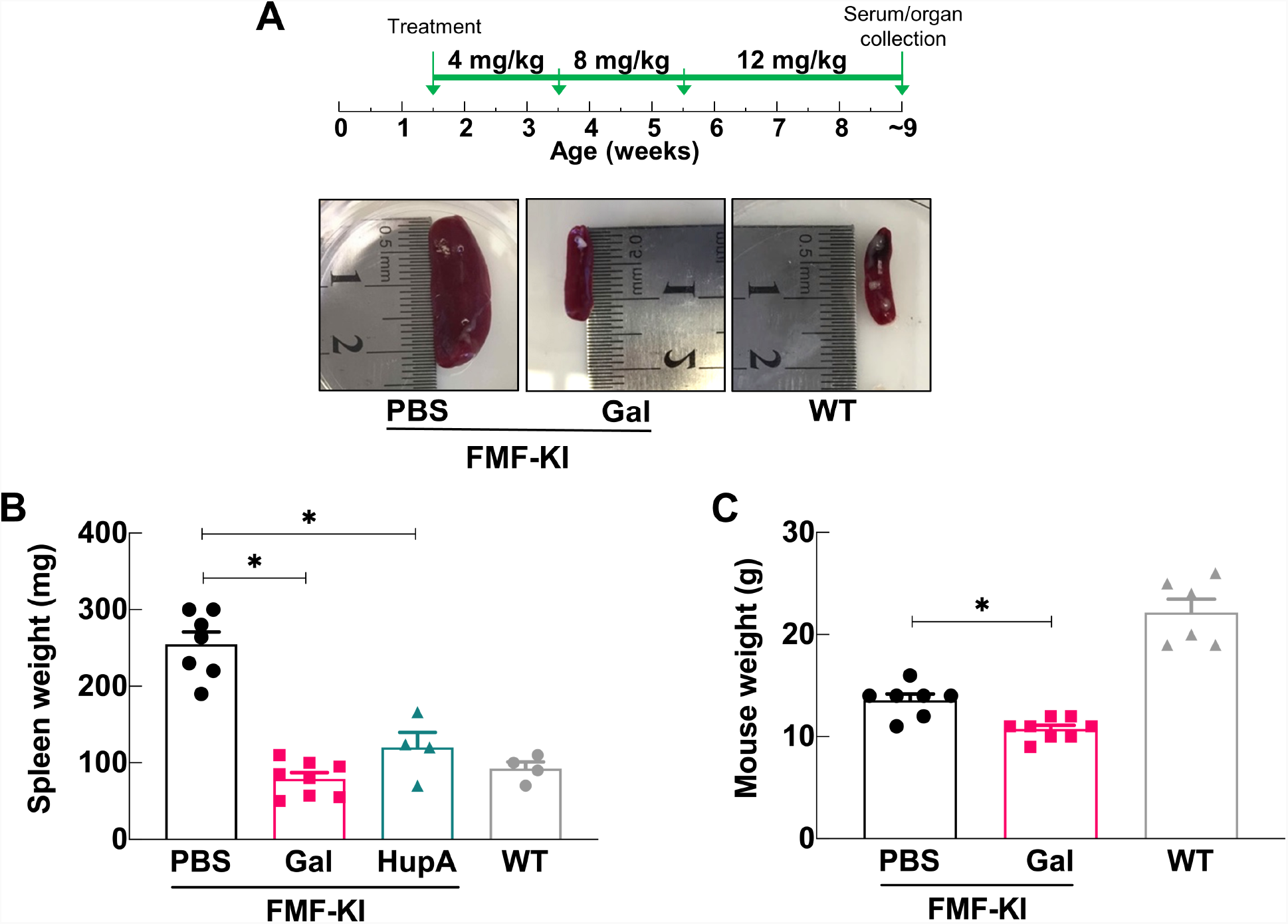
Galantamine prevents massive splenomegaly in chronically treated FMF-KI mice. **(A)** FMF-KI mice were treated with an escalated dose of galantamine (4-12 mg/kg) or PBS twice daily for ∼8 weeks starting at age 1.5 weeks (upper panel). The lower panels include representative images of FMF-KI mouse spleens harvested at the end of treatment compared with wildtype (WT) spleen (weights: PBS = 330 mg; gal = 70 mg; WT = 90 mg). **(B)** Spleen weights (mg) from galantamine-, huperzine A-, and PBS-treated FMF-KI mice compared with WT. FMF-KI mice were treated with huperzine A (1 mg/kg twice daily) for 4-5 weeks starting at the age of ∼4 weeks and compared to the control group in the galantamine vs. PBS experiment (first column). (Data presented as mean ± SEM; \**P* < 0.025 by Mann-Whitney *U* test with Bonferroni correction). **(C)** Body weights (g) of galantamine- and PBS-treated FMF-KI mice at the end of treatment period. In all figures, wild type measurements are included for comparison only.

### Galantamine inhibits leukocyte accumulation in sites of inflammation

FMF-KI mice exhibit an increase in myeloid cell numbers in multiple sites, including the spleen, skin, and peritoneal cavity, with a preponderance of Ly6G^+^ neutrophils relative to F4/80^+^ macrophages [19]. To examine whether the reduction in spleen size observed with long-term galantamine treatment affected leukocyte accumulation in inflamed tissues, we determined the number and composition of myeloid cells in the spleen and peritoneal cavity of galantamine- and PBS-treated FMF-KI mice. The total number of cells in the peritoneum and splenocytes in the spleen was significantly lower in galantamine-treated FMF-KI mice compared with PBS-treated controls (Fig 2 A & B, S 1 A). Despite this decrease in size and cellularity, flow cytometric analysis of splenocytes from galantamine-treated mice revealed no significant difference in the percentage of CD11b^+^ cells compared to controls, although one galantamine-treated animal exhibited a ∼40% reduction in the percentage of these cells (Fig 2 C). Further, examining the percentage of neutrophils relative to macrophages within CD11b^+^ cells revealed no significant difference between the two groups (Fig S1 B).

**Figure 2.**
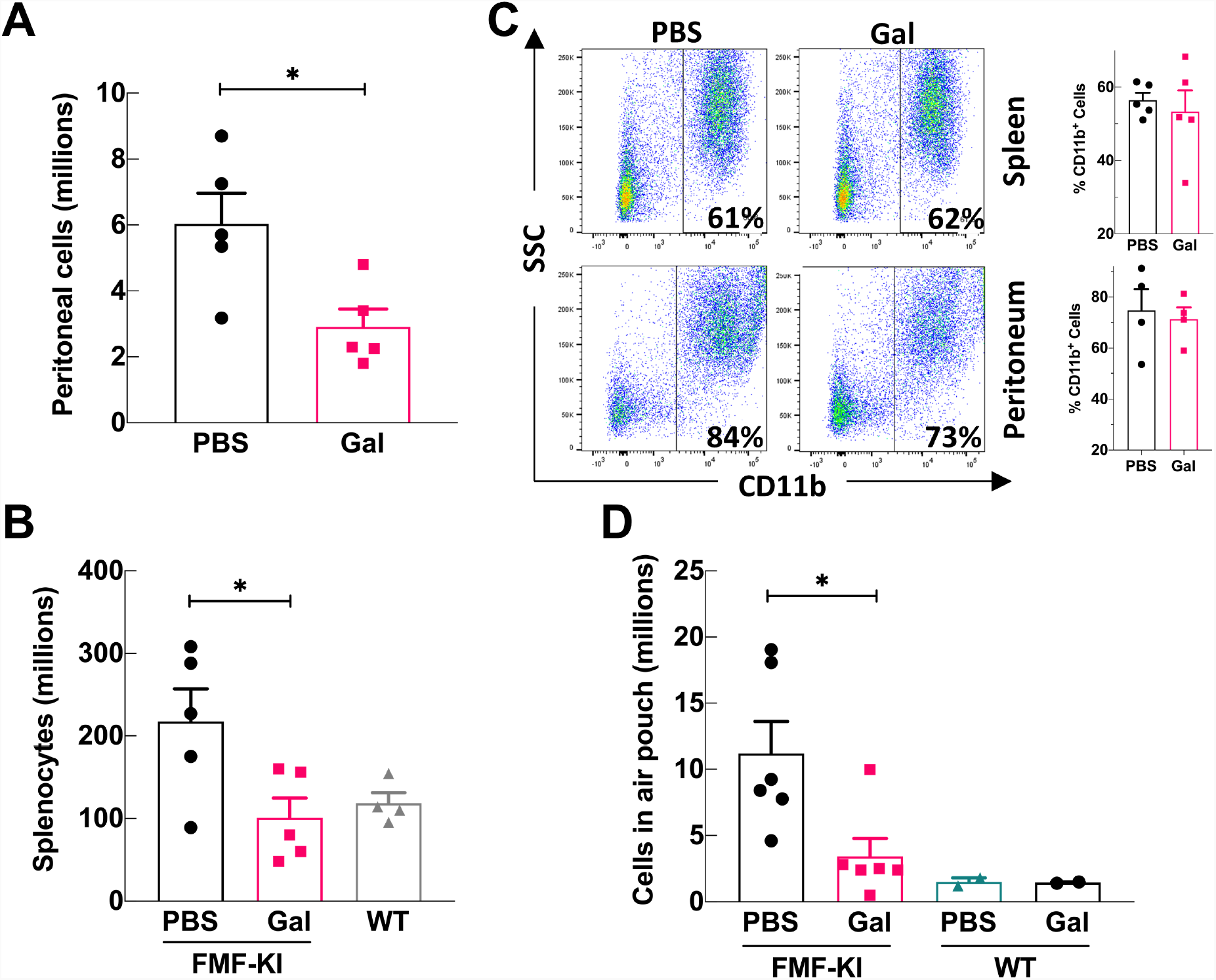
Galantamine-treatment inhibits leukocyte trafficking to inflammatory sites in FMF-KI mice. **(A and B)** Total number of cells in peritoneal washes (A) and total splenocyte counts (B) from galantamine- and PBS-treated FMF-KI mice at the end of 4-8 weeks of treatment (data presented as mean ± SEM; \**P* <0.5 by Mann-Whitney *U* test). **(C)** Example flow cytometric analysis of splenic and peritoneal cells from galantamine- and PBS-treated FMF-KI mice after 4-8 weeks of treatment. Total splenocytes were analyzed for relative frequencies of CD11b^+^. The percentage of CD11b^+^ cells from the spleens of galantamine- and PBS-treated FMF-KI mice (n = 5 in each group) is shown next to the flowcytometry panels. **(D)** Total number of cells in a subcutaneous dorsal air pouch from galantamine- and PBS-treated FMF-KI mice after 6 days of treatment (data presented as mean ± SEM; \**P* <0.05 by Mann-Whitney *U* test; WT is included for comparison only).

To determine whether effects on cellular accumulation were restricted to the spleen—as an organ directly modulated by the inflammatory reflex [23]—we decided to examine galantamine’s effects on systemic cell trafficking and inflammation using a subcutaneous air pouch model, which resembles a body cavity. After 6 days of treatment with galantamine (6-12 mg/kg, twice daily) or PBS, pouch contents were collected and analyzed. Treatment with galantamine significantly reduced the number of infiltrating cells in the air pouch compared with treatment with PBS (Fig 2 D). Notably, short-term treatment (∼1 week) with galantamine tended to reduce spleen size in treated air pouch mice, although not completely resolving the splenomegaly (Fig S1 C). Taken together, these data suggest that galantamine attenuates inflammatory cell infiltration into the spleen and body cavities.

**Fig. S1.**
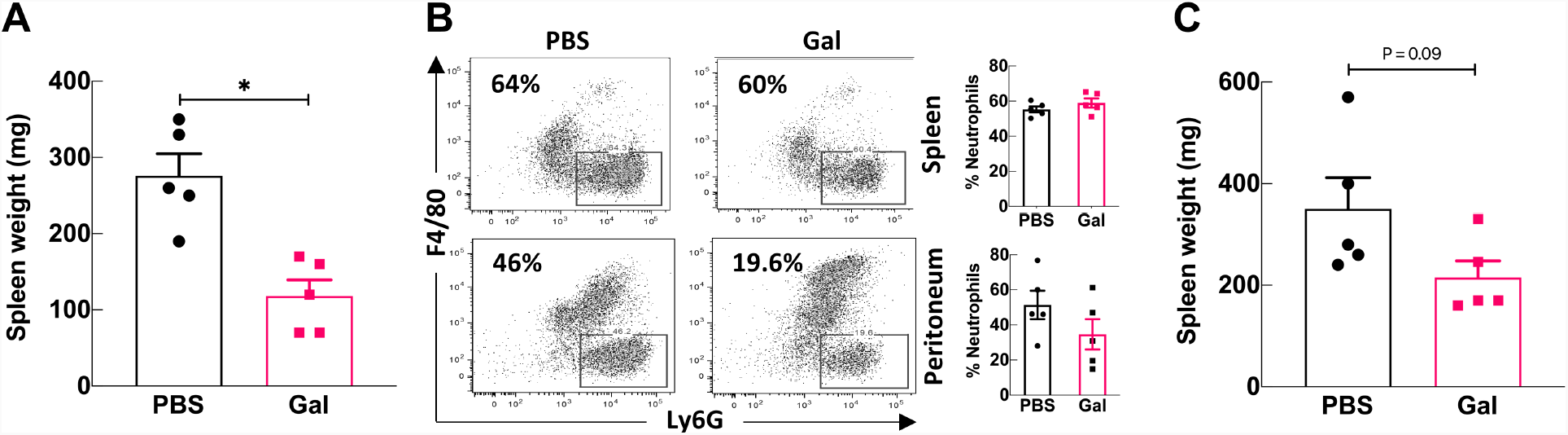
**(A)** Weights (mg) of FMF-KI spleens used for flow cytometric analysis after 4-8 weeks of treatement with galantamine or PBS. **(B)** Example flow cytometric analysis of splenic and peritoneal cells after 4-8 weeks of treatment. CD11b^+^ cells were analyzed for relative frequencies of Ly6G^+^ neutrophils and F4/80^+^ macrophages. The percentage of neutrophils in CD11b^+^ cells from the spleens and peritonea of galantamine- and PBS-treated FMF-KI mice (n = 5 in each group) is shown next to the flowcytometry panels. **(C)** Spleen weights (mg) from galantamine- and PBS-treated mice after 6 days of treatment with galantamine (twice daily, escalated to 12 mg/kg). Data presented as mean ± SEM, *P* by Mann-Whitney U test.

### Galantamine lowers markers of chronic inflammation

Uncontrolled inflammation in FMF patients leads to serum amyloid A (SAA) deposition in the kidneys and the development of anemia. To determine the effect of galantamine treatment on these two clinical markers of chronic inflammation, we measured renal amyloid deposition and complete blood cell counts in FMF-KI mice treated with either PBS or galantamine twice daily for 8 weeks. Long-term treatment with galantamine decreased SAA deposition in the kidneys, as well as the levels of SAA in the sera of FMF-KI mice (Fig. 3 *A*). It also restored mean values for hemoglobin concentration, hematocrit, and mean corpuscular volume to within or close to normal ranges [24], partially resolving the FMF-associated microcytic anemia (Fig. 3 *B*). No significant differences in blood neutrophil or lymphocyte counts were observed in galantamine-treated mice (data not shown).

**Figure 3.**
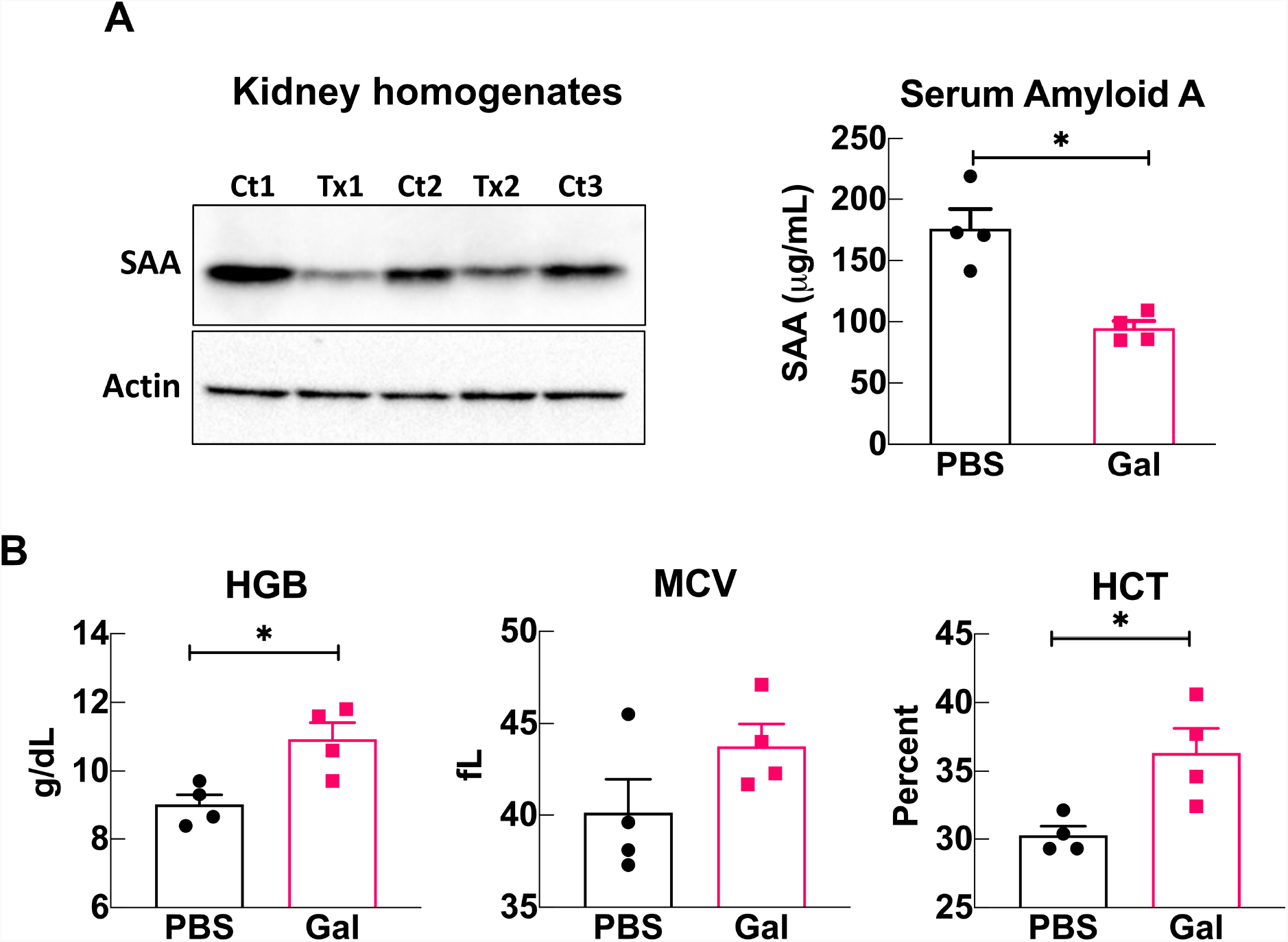
Galantamine lowers markers of chronic inflammation in FMF-KI mice. **(A)** Serum (right panel) and kidney deposited (left panel) SAA. Kidney homogenates from galantamine-treated (Tx) and PBS-treated (Ct) FMF-KI mice were probed for SAA deposition after 8 weeks of treatment. **(B)** Hemoglobin (HGB), mean corpuscular volume (MCV), and hematocrit (HCT) levels in galantamine- and PBS-treated FMF-KI mice at the end of 8 weeks of treatment (data presented as mean ± SEM; n = 4 per group; \**P<0*.*5* by Mann-Whitney *U* test).

The original study describing FMF-KI mice reported elevated levels of serum IL-1β in affected animals. In our study, IL-1β levels in both PBS- and galantamine-treated FMF-KI mice were near the limit of detection at all time-points tested; however, we found that IL-6 was significantly elevated in galantamine-treated mice compared to PBS-treated controls (Fig. S2 *A*). Serum levels of TNFα, IFNγ and IL-10 were low in both groups and no significant differences were observed (Fig S2 *B*). These data indicate that treatment with galantamine suppresses some of the key chronic inflammatory complications associated with FMF.

**Fig. S2.**
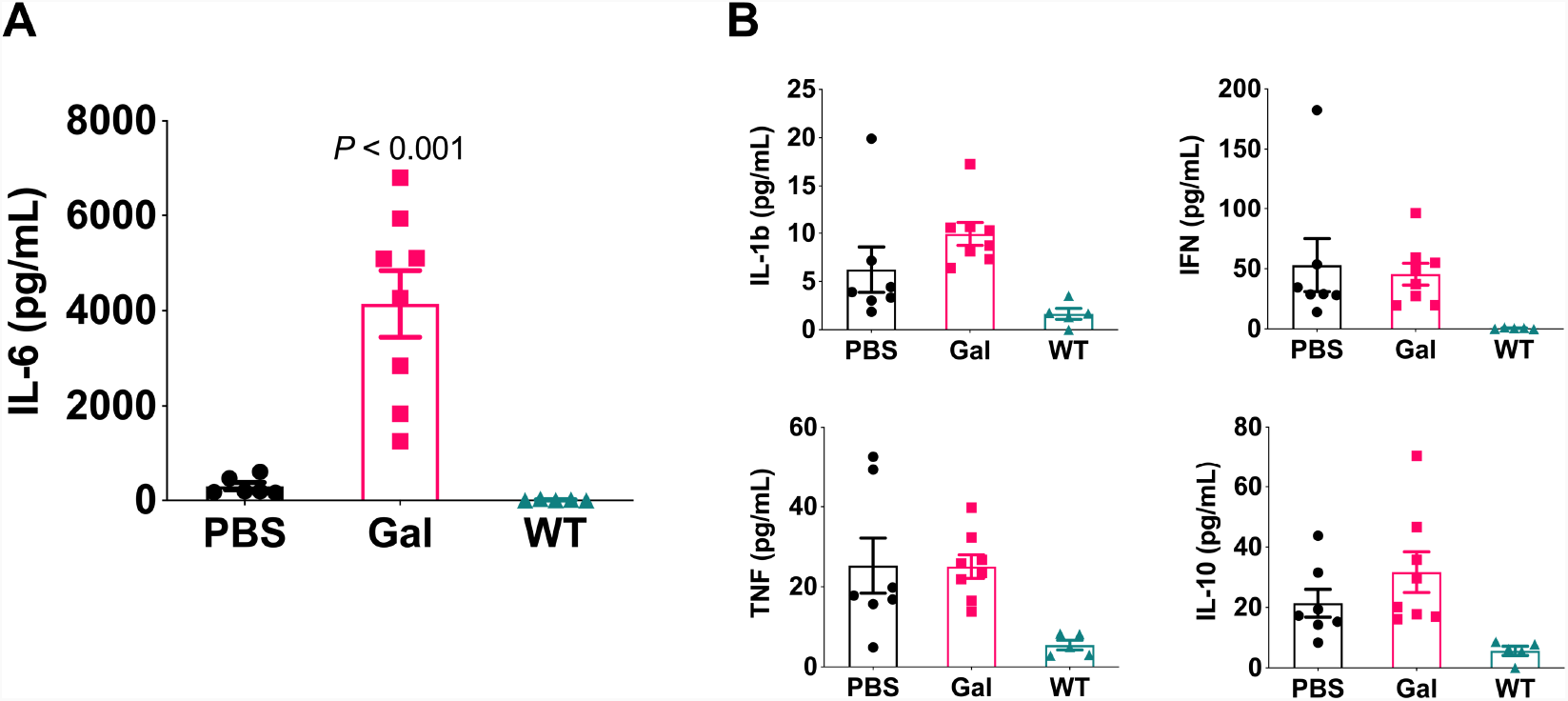
Serum cytokine levels from FMF-KI mice after 8 weeks of treatment with galantamine. **(A and B)** IL-6 (A), IL-1β, INF-γ, TNF-α, and IL-10 (B) levels in sera from FMF mice measured by multiplex ELISA (data presented as mean ± SEM, *P* value by Mann-Whitney U test compared to PBS).

### Galantamine controls splenomegaly in vagotomized and α7nAChR knockout (α7-KO) FMF-KI mice

Previously, several disease models have shown that galantamine required an intact vagus nerve and expression of the α7-nAChR to elicit its anti-inflammatory actions [10,11,25]. To determine whether galantamine mediates its anti-inflammatory effects in FMF-KI mice via this pathway, we tested whether an intact vagus nerve and α7-nAChR expression were required for the actions of galantamine on FMF-KI splenomegaly. In the first set of experiments, we severed the vagus nerve of FMF-KI mice at the age of 7 weeks, allowed the mice to recover for one week, and then initiated treatment with either PBS or galantamine. After 4 weeks of treatment, we found that galantamine-treated vagotomized mice had significantly smaller spleens than PBS-treated vagotomized controls (Fig. 4 *A*). We also compared the change in spleen dimensions within each mouse and found that all spleens in the galantamine-treated mice decreased in surface area, whereas spleens in the PBS-treated group increased (Fig. 4 *B*). In a second set of experiments, FMF-KI/α7nAChR-KO (FMF/α7-KO) mice, which had a similar disease phenotype to α7-intact FMF-KI mice including spleen size (Fig. 4 *C*), were treated with either PBS or galantamine for 4 weeks, and then spleen size and weight were evaluated. After 4 weeks of treatment with galantamine, α7nAChR deficient FMF-KI mice had significantly smaller spleens compared with PBS-treated controls (Fig. 4 *C*). To examine the role of cholinergic signaling via other receptors in FMF inflammation, we treated FMF macrophages with cholinergic agonists *ex vivo*. Treatment of LPS-primed FMF bone marrow-derived macrophages (BMDMs) with either Ach or nicotine failed to inhibit IL-1β release (Fig. S3 A) compared with the standard of care colchicine. These results indicate that galantamine’s effects on splenomegaly in FMF-KI mice do not require an intact vagus nerve or α7-nAChR signaling and suggest that cholinergic signaling might not be a direct effector mechanism on immune cells in this model.

**Figure 4.**
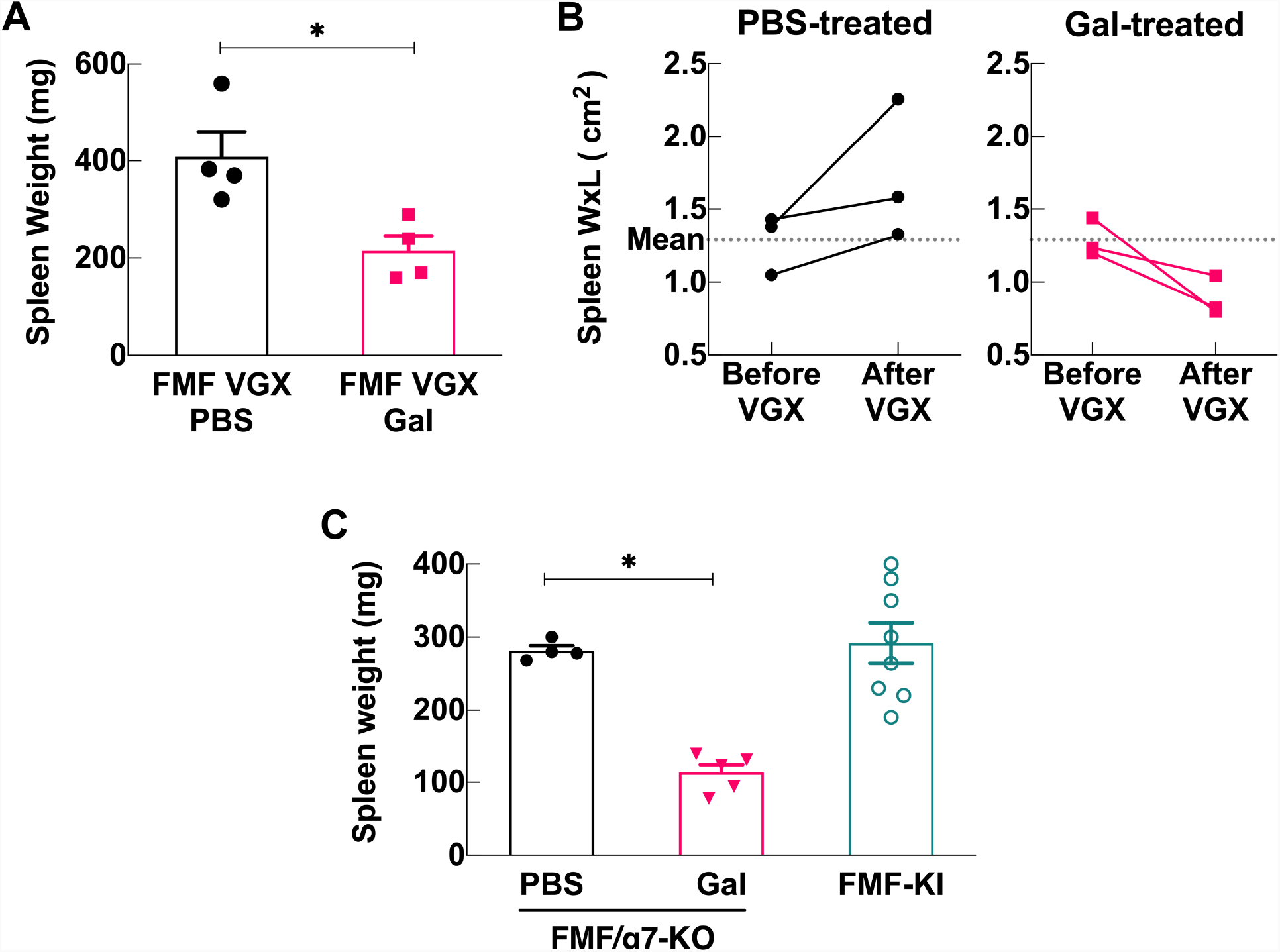
Galantamine controls splenomegaly in vagotomized (VGX) and α7nAChR knocked-out FMF-KI (FMF/α7-KO) mice. **(A)** Spleen weights (mg) from galantamine- and PBS-treated vagotomized mice after ∼4 weeks of treatment. **(B)** Spleen area (Width x Length cm^2^) before abdominal VGX (measured intra-operatively during VGX procedure) compared to area after 4 weeks of galantamine or PBS treatment. **(C)** Spleen sizes (mg) from galantamine- and PBS-treated α7nAChR deficient FMF-KI mice after 4-6 weeks of treatment with galantamine (escalated to 12 mg/kg twice daily) starting at age ∼4-5 weeks (treated n=5; control n=4). Spleen weights (mg) from α7-intact mice at ∼9-10 weeks of age (historical) are included for comparison only. Data presented as mean ± SEM; \**P <* 0.05 by Mann-Whitney *U* test with Bonferroni correction.

**Fig S 3.**
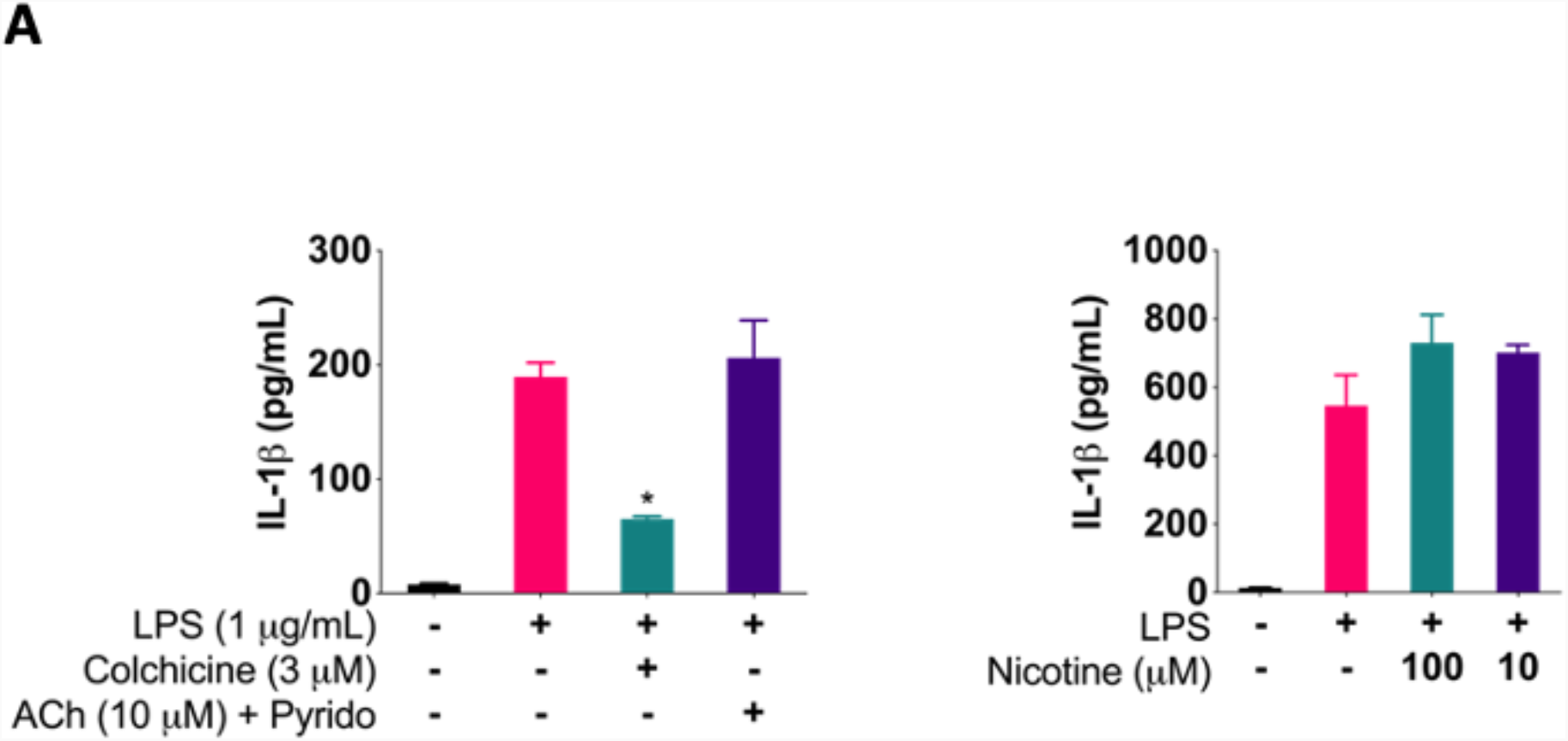
(A) IL-1β release measured by ELISA from FMF-KI BMDM primed with LPS and treated with Ach (with 100 mM pyridostigmine) and colchicine (left panel) or nicotine (right panel). Data presented as mean ± SD from duplicate measurement, \**P* <0.05 by ANOVA.

## Discussion

Galantamine has been used as a vagus nerve stimulator in various inflammatory models to inhibit cytokine production and prevent disease [18]. In this study, we utilize a mouse model of FMF to evaluate the therapeutic potential of galantamine in autoinflammation. Our findings demonstrate that long-term treatment of FMF-KI mice with galantamine attenuates several of the chronic signs and complications of the disease. This effect is likely due to inhibiting macrophage overactivation, based on the reported actions of galantamine in other models of inflammation.

FMF-KI mice exhibit dysregulated macrophage activation and IL-1β over-production, which serves to create a pro-inflammatory milieu that drives continuous neutrophil recruitment into inflamed tissues, causing splenomegaly, joint swelling, peritonitis, and hair loss [19]. Herein we demonstrate that long-term treatment with galantamine attenuates some of these inflammatory features and limits the number of inflammatory cells in affected tissues, including the spleen and a subcutaneous air pouch, suggesting that galantamine inhibits macrophage activation, thereby interrupting cytokine production and inflammatory cell recruitment. This notion is supported by previous work demonstrating phenotypic rescue of FMF-KI mice upon genetic deletion of the IL-1β receptor, which indicates that blocking the action of IL-1β ameliorates disease in this model [19]. Notably, however, despite showing a decrease in the total number of inflammatory cells in the spleen, we did not observe a significant change in the percentage of CD11b+ cells nor neutrophil to macrophage ratio, which remained high in almost all animals despite the resolution of splenomegaly. This residual inflammation most likely relates to galantamine’s mechanism of action, which might only induce transient suppression of the inflammatory cascade in FMF rather than complete blockade, especially as these mice harbor a constitutively active inflammasome and exhibit a severe form of the disease.

In addition to recruiting inflammatory cells, cytokines induce the production of SAA, an acute phase protein capable of depositing in tissues and causing amyloidosis. SAA aids in propagating inflammation and is produced mainly by the liver in response to circulating cytokines [26]. In FMF patients, SAA levels are used to monitor disease activity and response to treatment [27]. Persistently elevated cytokines can also lead to the development of anemia of chronic inflammation by mediating changes in iron metabolism and erythropoiesis as well as skewing hematopoietic stem cell differentiation towards myelopoiesis [28]. Treating FMF mice with galantamine lowered the levels of circulating SAA and the amount of amyloid deposited in kidneys as well as attenuated the anemia of chronic inflammation. A similar improvement in anemia was recently demonstrated in FMF-KI mice upon deletion of gasdermin D [29], a pore-forming protein required for IL-1β release. These findings further support the possibility that galantamine likely controls macrophage overproduction of cytokines thereby preventing their chronic sequalae.

Previous studies utilizing galantamine in inflammatory models have found that severing the abdominal vagus nerve or blocking the α7 nicotinic receptor abrogates the drug’s anti-inflammatory activity [10,11]. Here, we found that neither intervention affected the observed decrease in splenic size of galantamine-treated FMF-KI mice, suggesting the involvement of a different effector mechanism in this model. However, the fact that galantamine’s actions were replicated by another AchE Inhibitor (huperzine A) suggests that this mechanism still requires cholinergic signaling. It is likely that peripheral and/or central potentiation of cholinergic signaling with these AchE Inhibitors lead to the engagement of a different neuro-immune pathway and/or effector molecules that control inflammation. Future work will focus on identifying the circuits and molecules at play in this model. Interestingly, blocking the vagus or α7 receptor did not appear to worsen the FMF inflammatory phenotype unlike previous reports demonstrating increased inflammation with these interventions in acute endotoxemia. This might indicate an altered role of this pathway in lifelong severe inflammatory disease.

Despite the striking decrease in splenomegaly, galantamine treatment did not completely prevent hair loss and joint swelling—which appeared less severe in treated animals, but due to variability in the age of onset and severity of these manifestations, we were unable to quantify them reliably. We were also unable to detect the previously reported elevated levels of IL-1β in these animals. Beyond technical challenges, this could have several explanations. It could be due to the short half-life of IL-1β and its strong binding to membrane and serum receptors, which is most likely why FMF patients have surprisingly low levels of IL-1β during attacks [30]. Alternatively, we could have been sampling blood during disease remission thereby missing intermittent IL-1β spikes. This is supported by the observed low level of other inflammatory cytokines in our samples. Also, the IL-1β production mechanism could be susceptible to depletion in chronic disease or could vary with housing conditions, time of day, and age. This could be gleaned from the original report describing the model, in which serum IL-1β levels (mean value of approximately 300 pg/ml) exhibited considerable variability with most of the readings clustering below the mean [19]. Moreover, subsequent studies utilizing the FMF-KI mouse model reported lower levels of serum IL-1β compared with the original report, with readings clustering around 50 pg/ml [29,31]. Surprisingly, however, FMF-KI mice treated with galantamine had significantly higher levels of IL-6 compared with controls. While this could indicate active inflammation, IL-6 is known to be a pleiotropic cytokine with both pro- and anti-inflammatory actions based on the cellular and molecular milieu [32,33]. For example, IL-6 has been shown to upregulate IL-1 receptor antagonist and IL-10 while inhibiting NF-kB activation and promoting M2 macrophage polarization via IL-4 induction [34,35]. Therefore, IL-6 might be contributing to attenuation of inflammation in galantamine treated mice via a similar mechanism. Skeletal muscles could be another potential source of IL-6, which is released as a myokine in large amounts upon muscle contraction [36,37]. AchE inhibitors, such as galantamine, have been shown to cause muscle fasciculations, especially at higher doses, due to their actions on the neuromuscular junction [38], which would release IL-6 into the circulation. Nevertheless, the exact source and role of IL-6 in galantamine treated animals remain to be evaluated. Overall, even though we did not observe a direct effect of treatment on reducing levels of inflammatory cytokines, galantamine attenuated multiple chronic inflammatory markers known to be mediated by cytokines. Future work will focus on investigating the underlying mechanism of galantamine’s actions in this model.

In conclusion, our results demonstrate that galantamine attenuates several cytokine-mediated inflammatory markers and reduces renal amyloidosis in FMF-KI mice. These findings are of interest for developing galantamine as a novel treatment for FMF, an inflammasome-mediated chronic autoinflammatory disease, and possibly for other monogenic and complex autoinflammatory diseases.

## Acknowledgements

We would like to thank Christopher Colon (Flow Cytometry Core Facility at Feinstein Institutes) for providing training and assistance and Lionel Blanc (Laboratory of Developmental Erythropoiesis) for providing training and access to the hematology analyzer.

## Authors Contribution

I.T.M., Y.A. conceived and designed experiments; I.T.M., M.T., M.O. performed experiments; I.T.M., Y.A., B.S., B.D., V.A.P., J.J.C., D.L.K. analyzed and interpreted data; I.T.M. prepared figures; I.T.M. drafted manuscript; all authors revised and edited manuscript.

